# The earliest allopolyploidization in tracheophytes revealed by phylotranscriptomics and morphology of Selaginellaceae

**DOI:** 10.1101/2024.01.08.574748

**Authors:** Jong-Soo Kang, Ji-Gao Yu, Qiao-Ping Xiang, Xian-Chun Zhang

## Abstract

Selaginellaceae exhibit extraordinary evolutionary history in which they survived and thrived during the Permian–Triassic extinction and did not undergo polyploidization. Here, we reconstructed the phylogenetic relationships of Selaginellaceae by applying large-scale nuclear genes from RNA-seq, and found that each group showed phylogenetic incongruences among single-gene trees with different frequencies. In particular, three different phylogenetic positions of the *sanguinolenta* group were recovered by different nuclear gene sets. We evaluated the factors that might lead to the phylogenetic incongruence of the *sanguinolenta* group and concluded that hybridization between each ancestor of two superclades is the most likely cause. We presented the supporting evidence from gene flow test, species network inference, and plastome-based phylogeny. Furthermore, morphological characters and chromosomal evidence also lend support to the hybrid origin of this group. The divergence time estimations, using two gene sets respectively, indicated the splits between the *sanguinolenta* group and each related superclade happened around the same period, implying that the hybridization event probably occurred during the Early Triassic. This study reveals an ancient allopolyploidization with integrative evidence and robust analyses, which sheds new light on the recalcitrant phylogenetic problem of the *sanguinolenta* group and reports the polyploidization in the basal vascular plants, Selaginellaceae.

## Introduction

Lycophytes share a common ancestry with other tracheophytes that dates back nearly 400 million years, making them a key group to study the evolution of land plants. Selaginellaceae is the largest family of lycophytes, consisting of a single genus, with approximately 750 species, distributed globally in the northern and southern hemispheres (Jermy, 1990; Zhang et al., 2013). This family has been attracting researchers’ attention with distinctive genome evolution such as lacking whole-genome duplication (WGD) event, unconventional organelle genomes, and environmental adaptation through desiccation-tolerant species (Banks et al., 2011; Jiao et al., 2011; Li et al., 2015; Baniaga et al., 2016; VanBuren et al., 2018; Xu et al., 2018). Selaginellaceae is heterosporous, producing mega- and microspores in different sporangia, and has a special feature called rootlike rhizophores (Jermy, 1990; Zhou and Zhang, 2015; Weststrand and Korall, 2016a). Species in this family exhibit diverse variations in the position of rhizophores, the morphology of vegetative leaves and sporophylls, and the leaf arrangement (Jermy, 1990; Zhou and Zhang, 2015; Weststrand and Korall, 2016a).

Since DNA sequencing was applied in phylogenetics, phylogenetic incongruence between different gene trees has been frequently raised as a problem, which is generally interpreted by four factors such as gene duplication and loss, hybridization, gene transfer and introgression, and incomplete lineage sorting (ILS) (Maddison, 1997). It is challenging to differentiate them in case studies, especially when the events happened in the ancient period. Recently, progress has been achieved through research efforts, technical improvements, and empirical studies (Goulet et al., 2017; Folk et al., 2018; Kubatko and Chifman, 2019). The phylogenetic incongruence has also emerged as a recalcitrant problem in the carboniferous originated Selaginellaceae, particularly in the *Selaginella sinensis* and *S. sanguinolenta* groups (Korall and Kenrick, 2004; Zhou et al., 2015a; Weststrand and Korall, 2016b). According to the latest infrageneric classification, *Selaginella* consists of seven subgenera: *Selaginella*, *Rupestae*, *Lepidophyllae*, *Gymnogynum*, *Exaltatae*, *Ericetorum*, and *Stachygynandrum* (Weststrand and Korall, 2016a). The two problematic groups morphologically belong to subgenus *Stachygynandrum*, however their positions differed by different gene trees (Korall and Kenrick, 2004; Weststrand and Korall, 2016b) (Text S1).

Transcriptome (RNA-seq) has been demonstrated as an efficient approach to generating large-scale nuclear data (Hittinger et al., 2010; Wen et al., 2013; Huang et al., 2015). It provides large-scale nuclear genes for phylogenomic study, allowing for the successful resolution of deep phylogenetic relationships of major clades with significant statistical support (Wickett et al., 2014; Zeng et al., 2014; Huang et al., 2015, 2016; Xiang et al., 2017; Qi et al., 2018; Ran et al., 2018; Shen et al., 2018; Zhang et al., 2022; Chen et al., 2023). Given its potential, we conducted a deep phylogenomic reconstruction of Selaginellaceae based on RNA-seq data. The specific goal of this study was to provide the species network essentially to understand the phylogenetic relationships among subgenera and to explore the evolutionary history resulting in the enigmatic groups in this evolutionary key family.

## Results

### Phylogenetic reconstruction of Selaginellaceae

More than 3,000 single/low copy nuclear gene groups were used to select the orthologous genes. A total of 347 orthologous genes were eventually obtained for further analyses; the information in detail is described in supporting information (Text S2). According to the three positions of the *sanguinolenta* group, we divided the selected orthologous genes into three gene sets (Figure 1A). Three different phylogenetic positions of the *sanguinolenta* group are as follow: 1) sister to superclade A, including the four subgenera; 2) sister to superclade C, including subgenus *Stachygynandrum*; and 3) sister to the clade composed of superclades A and C (Figures 1A, 2A). The second and third positions of the *sanguinolenta* group, inferred from gene sets A and C respectively, were previously reported (Zhou et al., 2015a; Weststrand and Korall, 2016b; Du et al., 2020; Zhang et al., 2020; Chen et al., 2022; Tang et al., 2023). However, the position inferred from gene set B was newly revealed in this study (Figure 1B). The three positions inferred from three different gene sets were strongly supported by both concatenated ML and coalescent ASTRAL methods (Figures 1A, 2B). Using the combined three gene sets, the topologies inferred from both ML and ASTRAL methods were consistent with the topology using gene set C (Figure 2B). While the position of the *sanguinolenta* group was 100% supported in the concatenated ML method, the local posterior probability of the position was 38.8% in the coalescent ASTRAL method (Figure 2B).

**Figure 1.**
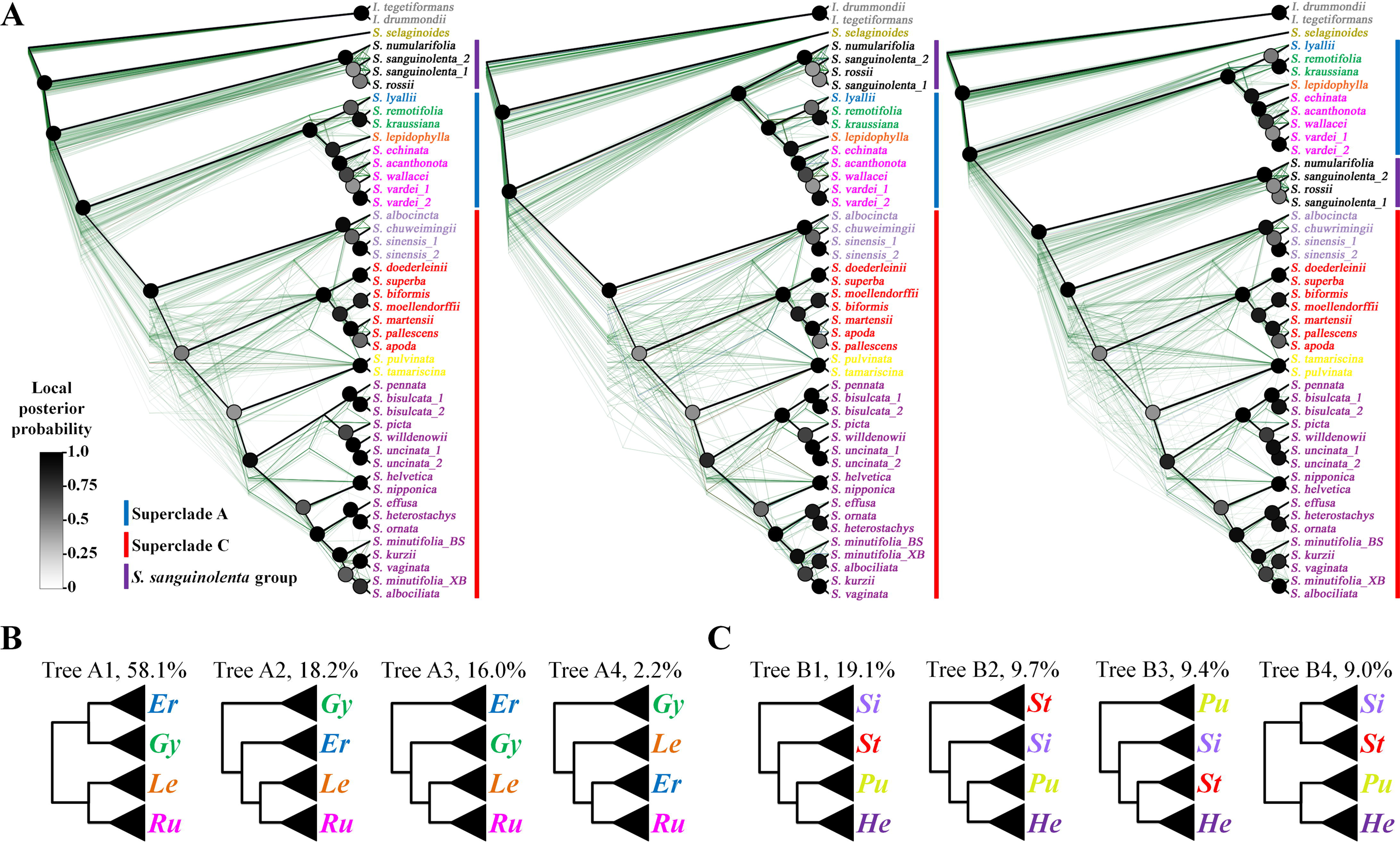
Coalescent-based nuclear phylogenies and topology frequencies for each group in Selaginellaceae. (A) Cladograms of the coalescent-based species tree (black line) and single gene trees for 130, 81, 136 gene sets (green lines) corresponding to the position of the *sanguinolenta* group. The node color represents local posterior support values calculated by ASTRAL, according to the scale shown on the left. (B) The most common topologies in single gene trees with the percentage of the topology. Er: subg. *Ericetorum*, Gy: subg. *Gymnogynum*, Le: sug. *Lepidophyllae*, Ru: subg. *Rupestrae*, Sa: the *sanguinolenta* group, Si: the *sinensis* group, St: subg. *Stachygynandrum sensu* Zhou and Zhang (2015), Pu: the *pulvinata* group, He: subg. *Heterostachys sensu* Zhou and Zhang (2015).

**Figure 2.**
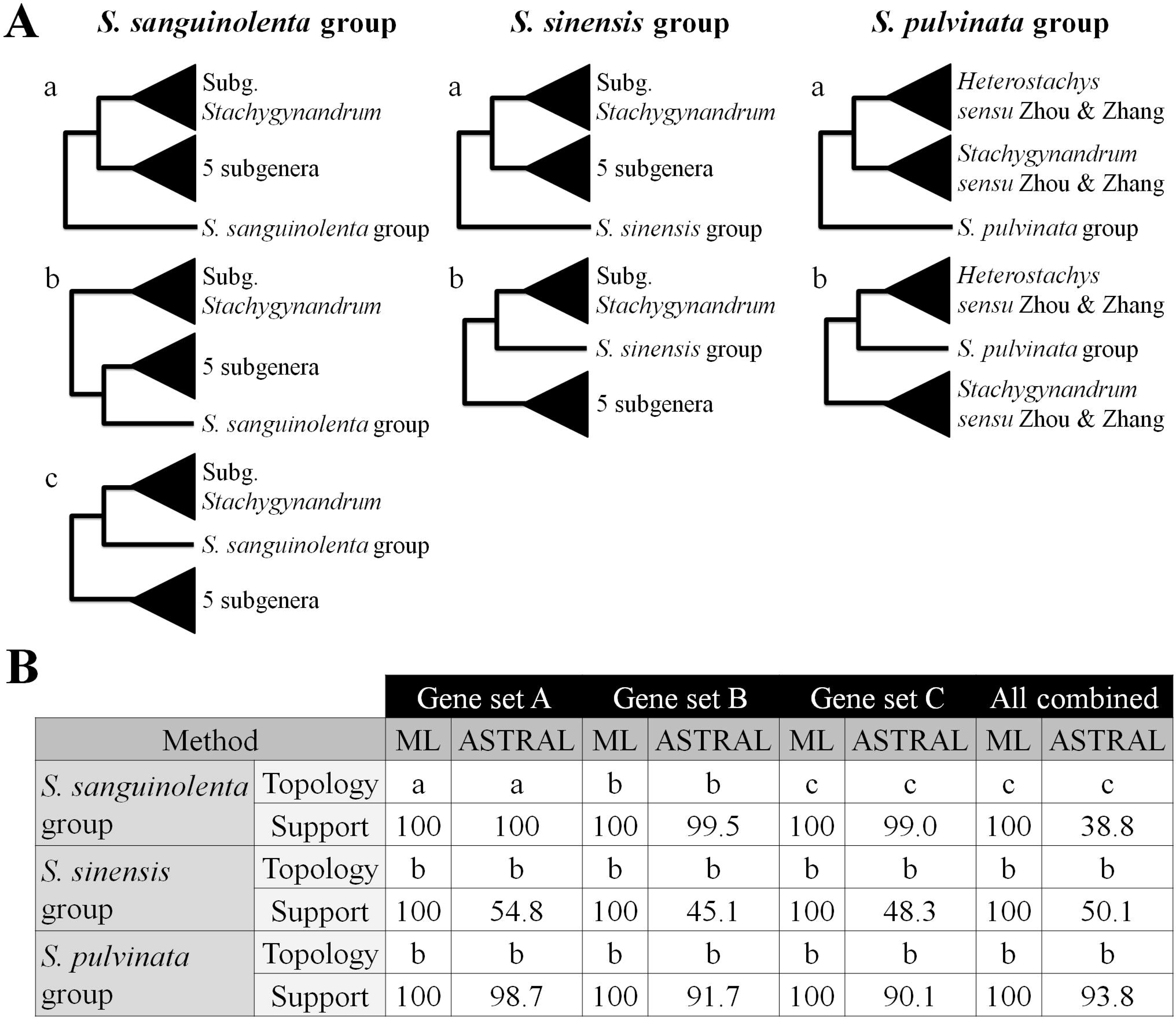
Summary of different phylogenetic relationships corresponding to the three enigmatic groups in Selaginellaceae. (A) Simplified topologies showing different phylogenetic relationships of three different groups. (B) Bootstrap support values and local posterior probabilities of each topology from concatenated ML and coalescent methods. The three groups, *S. sanguinolenta*, *S. sinensis*, and *S. pulvinata* groups are marked as Sa, Si, and Pu, in Figure 1.

Except for the position of the *sanguinolenta* group, the phylogenetic results of the three gene sets showed a consistent topology inferred from concatenated ML and coalescent ASTRAL methods with strong support values (Figure 1). *S. selaginoides*, representing subgenus *Selaginella*, was resolved in the most basal clade, sister to the all-rhizophoric *Selaginella*, with 100% support in all phylogenetic analyses (Figure 1A). The superclade A is composed of four subgenera, *Rupestrae*, *Lepidophyllae*, *Gymnogynum*, and *Ericetorum*, and the superclade C is composed of subgenus *Stachygynandrum sensu* Weststrand and Korall (2016a), including the enigmatic *sinensis* group (Figure 1A). Within the superclade A, both concatenated ML and coalescent ASTRAL analyses indicated that subgenera *Ericetorum* and *Gymnogynum* formed a subclade, and subgenera *Lepidophyllae* and *Rupestrae* formed the other subclade (Figure 1A). This topology was the most frequent one resulted from the 347 single genes (Figure 1B). Within the superclade B, both analyzing methods put the *sinensis* group as the basal group, and the *pulvinata* group (= subg. *Pulviniella sensu* Zhou and Zhang) was sister to the *Heterostachys* group (= subg. *Heterostachys sensu* Zhou and Zhang) (Figure 1A). This topology was the most frequent in the 347 single-gene trees (ca. 20% frequency), and three other topologies were about 10% frequency for each (Figure 1C). The monophyly of every subgenera and groups of subg. *Stachygynandrum* were well recovered with strong supports in all phylogenetic reconstructions (Figure 1A). According to the classification of Zhou and Zhang (2015), the *Heterostachys* group consists of five sections, four of which were included in this study. Among the four sections, sections *Homostachys* (*S. nipponica* and *S. helvetica*) and *Oligomacrosporangiatae* (*S. uncinata*, *S. willdenowii*, *S. picta*, *S. bisulcata*, and *S. pennata*) were monophyletic with strong supports, whereas sect. *Heterostachys* (*S. kurzii*) was nested within sect. *Tetragonostachyae* (Figure 1A). In the *Stachygynandrum* group (= subg. *Stachygynandrum sensu* Zhou and Zhang), five of seven sections were included in this study. Sect. *Ascendentes* (*S. doederleinii* and *S. superba*) and sect. *Pallescentes* (*S. pallescens* and *S. apoda*) formed their own clades; however, the monophyly of the other three sections could not be evaluated due to the single sampling (Figure 1).

### Intergroup gene flow and species network inference

Since the *sanguinolenta* group was monophyletic and sister to one or both of the two superclades (Figure 1), we defined three groups in the Selaginellaceae to test the gene flow and to infer the species network. The three clearly defined groups are as follow: 1) Group A for the superclade A, 2) Group B for the *sanguinolenta* group, and 3) Group C for the superclade C (Figure 3A). The three groups were used as ingroups, and *S. selaginoides* (subg. *Selaginella*) and two *Isoetes* species were defined as outgroups for the HyDe test (Blischak et al., 2018). A total of 444,491 nucleotides were used for this gene flow test. As a result, a significant hybridization signal was detected in the *sanguinolenta* group (Z-score=18.8572, P-value=∼0.0, γ=0.4511) (Figures 3B, C). Of 1000 bootstrap replicates, 96.3% detected the hybridization in the *sanguinolenta* group, and the estimated γ values (the inheritance probability from an ancestor of group A in this study) ranged from 0.0991 to 0.6559. The species network inference, using PhyloNet (Than et al., 2008), suggested that the *sanguinolenta* group (group B) was hybrid originated, and the potential parents were the common ancestors of the two superclades, respectively (Figure 4A). The estimated inheritance probabilities (γ) indicated that the genetic composition of the *sanguinolenta* group was 47.6% from the common ancestor of the superclade A (γ; group A), and the remaining 52.3% from the common ancestor of the superclade C (1-γ; group C) (Figure 4A). These results were consistent with the result from the hybridization detection using HyDe, as well as the inheritance probabilities (Figures 3B, C).

**Figure 3.**
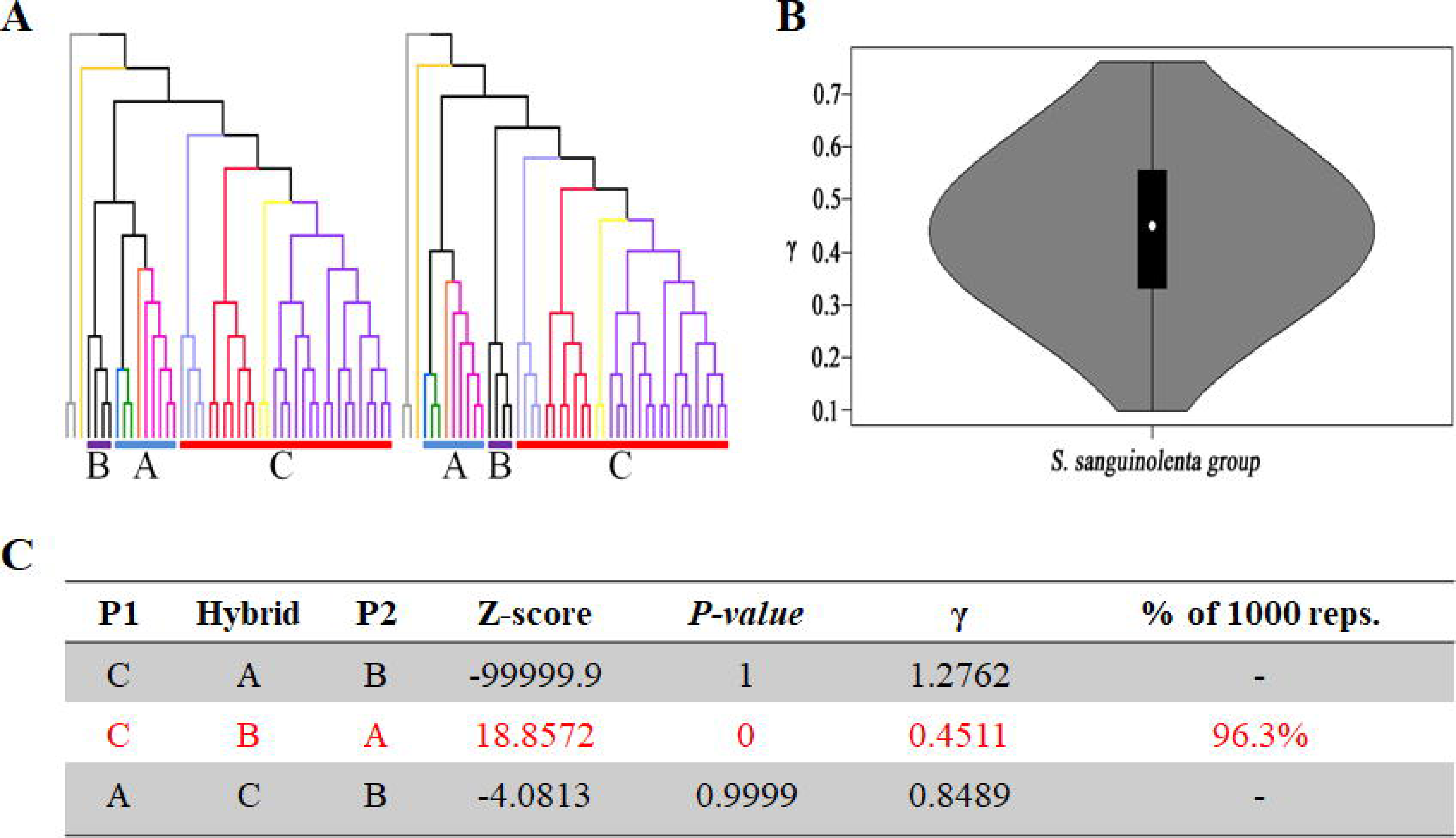
Summary of intergroup gene flow test using HyDe. (A) Designated three groups for the gene flow test. The superclade A (including four subgenera), the *sanguinolenta* group, and the superclade C (subgenus *Stachygynandrum*) are marked as groups A, B, and C, respectively. (B) Violin plot of distribution of γ calculated by the HyDe across 1000 bootstrap replicates. γ and 1-γ indicate inheritance probabilities from the putative parental groups A and C, respectively. (C) Test results of detecting hybridization among the three groups.

**Figure 4.**
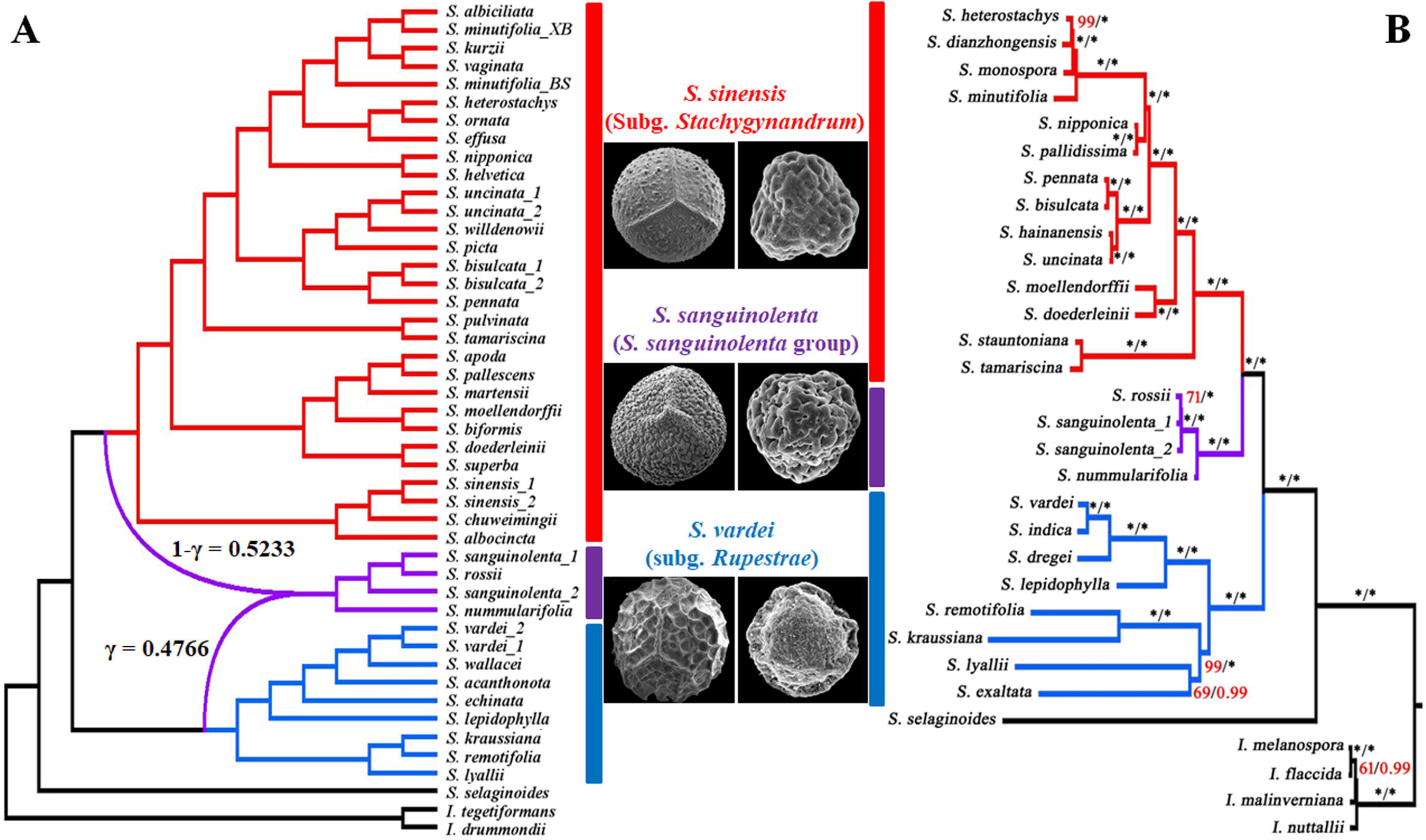
Species network inference from nuclear phylogenies and plastome-based phylogeny of Selaginellaceae. (A) Species network inferred from PhyloNet. (B) A plasome-based phylogeny adapted from Zhang, et al. (2020). The γ and 1-γ values indicate inheritance probabilities from the common ancestors of superclade A (blue) and superclade C (red), respectively. The values near the nodes are bootstrap and posterior probabilities from the ML and BI analyses, respectively. Asterisks indicate 100% supports, whereas the values lower than 100% support are shown in red. The megaspore (left) and microspore (right) of representative species for each group are presented in the middle of the species network inference (A) and the plastome-based phylogeny (B).

### Morphological and cytological evidences

To find morphological and chromosomal traits of the hybridization between the common ancestors of the two superclades, creating the *sanguinolenta* group, morphological and cytological characters were compared among subgenera or groups in Selaginellaceae (Table S1). The leaf arrangements evolved from spirally arranged to four-rows of vegetative leaves and sporophylls, and the leaf morphology evolved from monomorphic to dimorphic in shape for vegetative leaves and sporophylls (Table S1). Subg. *Selaginella*, the most basal group of the genus, possesses several ancestral characters, including helical arrangements of vegetative leaves and sporophylls, and the absence of rhizophores (Table S1). The position of the rhizophore is a key character to distinguish the two superclades. Species in the superclade A have the rhizophores on the dorsal side, whereas species in the superclade C have them on the ventral side (Table S1). Although the *sanguinolenta* group is morphologically similar to the superclade C, the rhizophore position of the *sanguinolenta* group is distinct from other species in the superclade C because the *sanguinolenta* group has the rhizophores on the dorsal side (Figure S1), which is the typical character of the superclade A.

In the spore morphology, the two superclades can be distinguished by the surface sculptures of megaspores. Species in the superclade A generally have a reticulate surface of megaspores, whereas species in the superclade C generally have a verrucate surface of megaspores (Figure S2; Table S1). Interestingly, the megaspore surface of the *sanguinolenta* group was closer to an intermediate form between reticulate of the superclade A and verrucate of the superclade C, and the microspore surface of the *sanguinolenta* group also showed an intermediate form between the two superclades (Figure S2; Table S1). Moreover, the chromosome number of the *sanguinolenta* group differed from species in the two superclades. While there are 18 or 20 chromosomes in most *Selaginella* species (Takamiya, 1993), species in the *sanguinolenta* group has a total of 30 chromosomes (Figure S3; Table S1). The 30 chromosomes were consistent in different species of the *sanguinolenta* group, e.g., *S. rossii* collected from Japan (Takamiya, 1993) and *S. sanguinolenta* collected from China (this study; Figure S3), suggesting that the 30 chromosomes are probably the synapomorphy of the whole group.

### Molecular dating for the ancient hybridization

Based on the species inference that the *sanguinolenta* group was originated by the hybridization between each common ancestor of superclades A and C, respectively (Figure 3, 4), we used two gene sets, B and C, in which the *sanguinolenta* group was resolved as sister to the superclades A and C, respectively (Figure 1, 3). Using penalized likelihood (PL) and MCMCtree methods, estimated divergence times based on two different gene sets (B and C) were shown in Figure S4. Divergence times inferred by the PL method were slightly earlier than those inferred by the MCMCtree method (Figure S4). Between the estimated divergence times by two methods, the difference between the results of two gene sets was greater in the PL method (Figure S4A), whereas that of two gene sets was almost consistent in MCMCtree (Figure S4B). Furthermore, a mega-fossil of *Selaginella anasazia* (Ash, 1972) found in the Late Triassic has been regarded as an ancestral species of subg. *Gymnogynum*, and the minimum age of the fossil was set at 210 Ma (Arrigo et al., 2013). The fossil of *S. anasazia* has dimorphic and decussate vegetative leaves as well as monomorphic sporophylls, which correspond to the two subgenera in this study, *Gymnogynum* and *Ericetorum* (Table S1), but the divergence times inferred by the PL method were much younger than the fossil’s minimum age (Figure S4A). Therefore, we eventually applied the MCMCtree method for estimating the ancient hybridization time creating the *sanguinolenta* group (Figure 5A).

**Figure 5.**
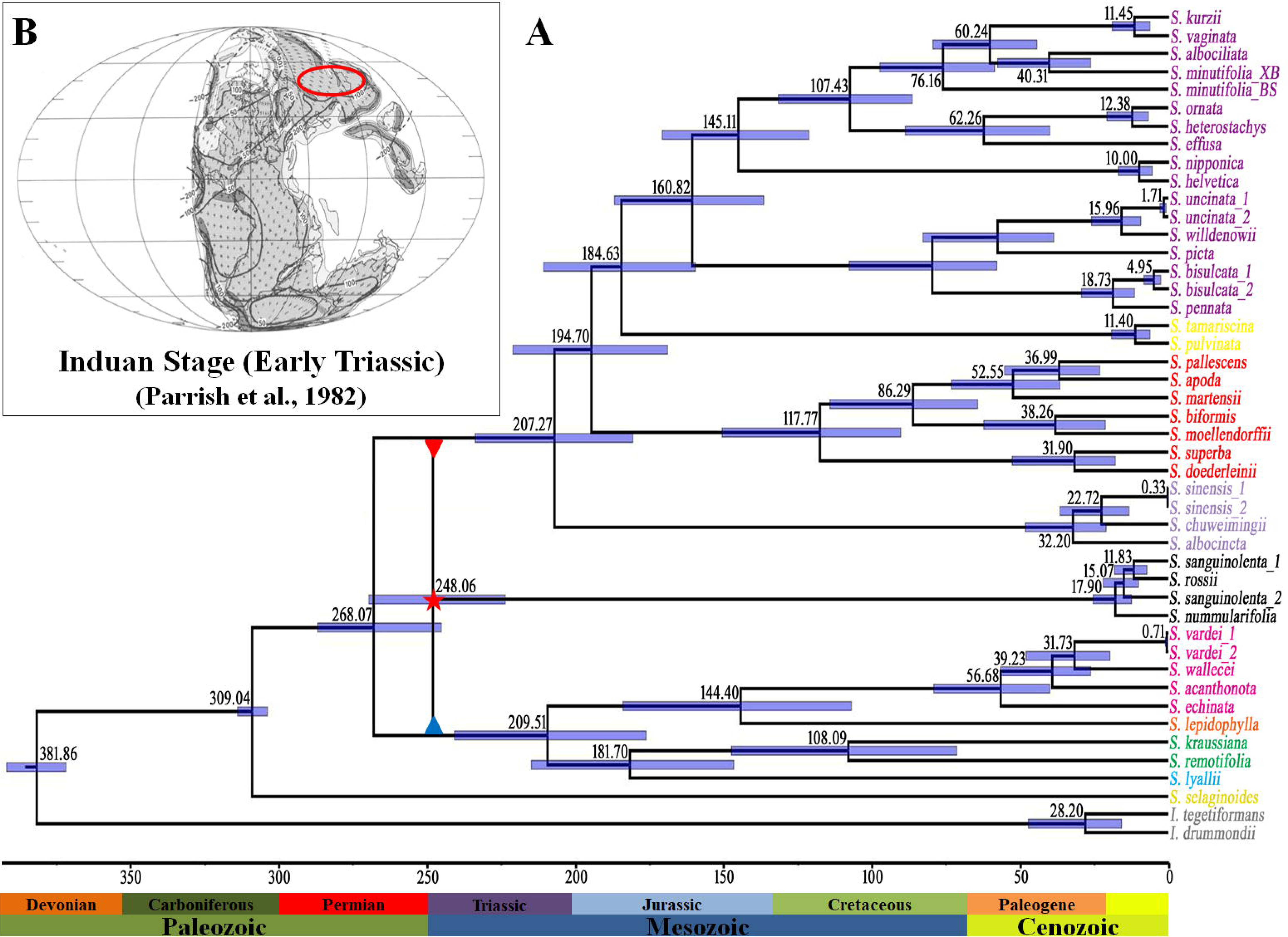
Time estimation of the hybridization between the two superclades in the Selaginellaceae creating the *sanguinolenta* group. (A) Divergence times for Selaginellaceae and the estimated time for the ancient hybridization that created the *sanguinolenta* group. The red star indicates the hybridization between the common ancestors of each superclade. The backbone of divergence time was inferred from gene set C using the MCMCtree method. (B) Predicted rainfall distribution of the Induan Stage in the Earliest Triassic by Parrish et al. (1982). The red circle indicates the current northern parts of China and some Siberian regions. The rainfall is divided into four categories: <50 = low rainfall, 50 to 100 = moderately low rainfall, 100 to 200 = moderately high rainfall, and >200 = high rainfall.

The divergence times of the two different data sets inferred by MCMCtree indicated that the split between the two superclades occurred at ca. 268 Ma (Figure S4B). Except for the *sanguinolenta* group, the divergence time estimations using the two different data sets indicated that the first split within the superclade A occurred at 196 Ma and 209 Ma, and the first split within the superclade C occurred at 217 Ma and 207 Ma, respectively (Figure S4B). According to the phylogenetic results and species network, the putative parents for the ancient hybridization were each common ancestor of the two superclades, respectively (Figures 1, 4). Conincidently, the estimated divergence times based on the two different gene sets using MCMCtree method indicated that the split times between the *sanguinolenta* group and each superclade were consistent, although the topologies were different, as sister to superclade A in gene set B and sister to superclade C in gene set C (Figures 1, 4, and S4). The divergence time between the *sanguinolenta* group and superclade A was around 247 Ma (268 to 223 Ma in 95%), and that between the *sanguinolenta* group and superclade C was around 248 Ma (269 to 223 Ma in 95%) (Figure S4B). Therefore, the hybridization time yielding the *sanguinolenta* group was probably ca. 248 Ma (269 to 223 Ma) during the Late Permian to Late Triassic (Figure 5A).

## Discussion

### Phylogenetic relationships within Selaginellaceae

Phylogenetic relationships within Selaginellaceae are still unclear, which is an early-diverged lineage of land plants. The relationships among subgenera differed in each single gene tree based on the plastid *rbcL*, 26S nrDNA, and two nuclear genes, *pgiC* and *SQD1* (Korall and Kenrick, 2004; Zhou et al., 2015a; Weststrand and Korall, 2016b). The confliction among subgenera was not settled, even though plastid and mitochondrial genome data, which are uniparentally inherited, were applied for the phylogenetic reconstructions. Recently, three plastome-based and one mitogenome-based phylogenetic studies have attempted to resolve the infrageneric relationships of this family, but failed due to the phylogenetic incongruences within superclades A (four subgenera) and C (subgenus *Stachygynandrum*), especially within superclade C (Du et al., 2020; Zhang et al., 2020; Zhou et al., 2022; Tang et al., 2023) (Figure 1A). Within the superclade A, the sister relationship between subgenera *Rupestrae* and *Lepidophyllae* was strongly supported by plastome- and mitogenome-based phylogenies (Du et al., 2020; Zhang et al., 2020; Zhou et al., 2022; Tang et al., 2023), but the phylogenetic relationships among subgenera *Gymnogynum*, *Ericetorum*, and *Exatatae* differed by data partitioning strategies (Du et al., 2020). Although failed to include representative species of subgenus *Exaltatae*, we confirmed the close relationships between subgenera *Rupestrae* and *Lepidophyllae* (over 90% in Figure 1B), between subgenera *Gymnogynum* and *Ericetorum* (most frequent in Figure 1B), respectively. We recovered three different topologies with high frequencies in the superclade A, where the species tree topologies show one most frequent topology (58.1%) and the two others with almost equal frequencies (18.2% and 16.0%) (Figure 1B), suggesting that the incongruences among subgenera of superclade A may be caused by the ILS, like those topology frequencies previously regarded as a signature of the ILS (Pamilo and Nei, 1988; Zhang et al., 2019) (Figure S5).

The phylogenetic relationships within the superclade C is more complicated than those of the superclade A. Similar to the *sanguinolenta* group, the phylogenetic position of the *sinensis* group remains unclear. The reported plastome from the *sinensis* group exhibited extraordinary genomic and genetic features, even when compared to other *Selaginella* species (Xiang et al., 2022), established as the most basal group of Selaginellaceae with a long branch (Zhou et al., 2022). In contrast, its mitochondrial genome features were similar to those of other *Selaginella* species, and the phylogenetic position of the *sinensis* group based on the mitogenomes was located within the superclade C (Tang et al., 2023). The mitogenome-based topology was about 9.7% of the total topology frequency revealed by transcriptome data in this study (Figure 1C). The most frequent position of the *sinensis* group is sister to the remaining species in the superclade C (Figure 1A, C), which well corresponds with the morphological characters, such as rhizophore position, leaf arrangement and morphology, and spore sculptures (Figure S1, S2; Table S1).

The phylogenetic position of the *pulvinata* group was resolved as sister group to the clade of the *Heterostachys* and *Stachygynandrum* groups by plastid phylogenies (Korall and Kenrick, 2004; Zhou et al., 2015a, 2022; Weststrand and Korall, 2016b; Du et al., 2020; Zhang et al., 2020; Chen et al., 2022; Zhang and Zhang, 2022). However, transcriptome-based results in this study did not consistent with the plastid phylogenies on the position of the *pulvinata* group (Figures 1, 2). Our results strongly indicated the *pulvinata* group and the *Heterostachys* group formed a monophyletic clade first, and then sister to the *Stachygynandrum* group (Figures 1, 2). This position was suggested by plasome-based phylogeny when considering RNA editing events (Du et al., 2020) and the mitogenome-based phylogeny (Tang et al., 2023). There are pervasive RNA editing in the Selaginellaceae organelle genomes to restore the accelerated mutations caused by the improper DNA repair system (Kang et al., 2020, 2022; Zhang et al., 2020; Xiang et al., 2022). The consensus position of the *pulvinata* group, achieved by the plastome-based phylogeny considering RNA editing, the mitogenome-based phylogeny, and the large-scale nuclear phylogenies, demonstrate the power of integrating the evidence from different genomes and the deep understanding of the data.

### Allopolyploidization of the *sanguinolenta* group

Hybridization with polyploidization, also known as allopolyploidization, is an important evolutionary force that can generate biological diversity, especially in plants (Stebbins, 1950; Grant, 1981; Soltis and Soltis, 2009; Goulet et al., 2017; Folk et al., 2018). It is challenging to elucidate the ancient hybridization because the genetic traits inherited from both parents should be investigated necessarily, but the traits diminish over time by mutation and recombination (Moody and Rieseberg, 2012; Folk et al., 2018). Four factors (gene duplication and loss, gene transfer and introgression, hybridization, and the ILS) are proposed to account for the widely discovered phylogenetic incongruences. Prior to discussing on the hybridization evidence, we discussed how we logically excluded the factors that cause phylogenetic incongruence step by step. Firstly, if gene duplication and loss were the factor causing the phylogenetic incongruence of the *sanguinolenta* group, we were able to detect and remove them during careful orthologous gene selection in this study. Secondly, if the phylogenetic incongruence was caused by gene transfer and introgression, one topology would be shown in most single gene trees, and the other topology would be shown rarely (Folk et al., 2018), but this was not the case in this family because the genes showing each topology appear to be common in the nuclear genome, such as 130, 81, and 136 genes for each topology (Figure 1A). Finally, we found evidence that phylogenetic incongruence did not result from the ILS. The first evidence was the split time difference of the two superclades (t*_i_* and t*_k_*, respectively, in Figure 6). If it was caused by ILS, the time difference between t*_i_* and t*_k_*should be greater (Sang and Zhong, 2000) (Figure S5), but they were similar in the Selaginellaceae (Figure 6). The other evidence was the topology frequencies among the three topologies (Figure 1). If it was caused by the ILS, one topology should be the most frequent, and the other two topologies were equally frequent among the three topologies (Pamilo and Nei, 1988) (Figure S5), but the number of single gene trees corresponding to the three topologies were 130, 81, and 136, respectively (Figure 1). One is the least frequent, the other two are almost equal.

**Figure 6.**
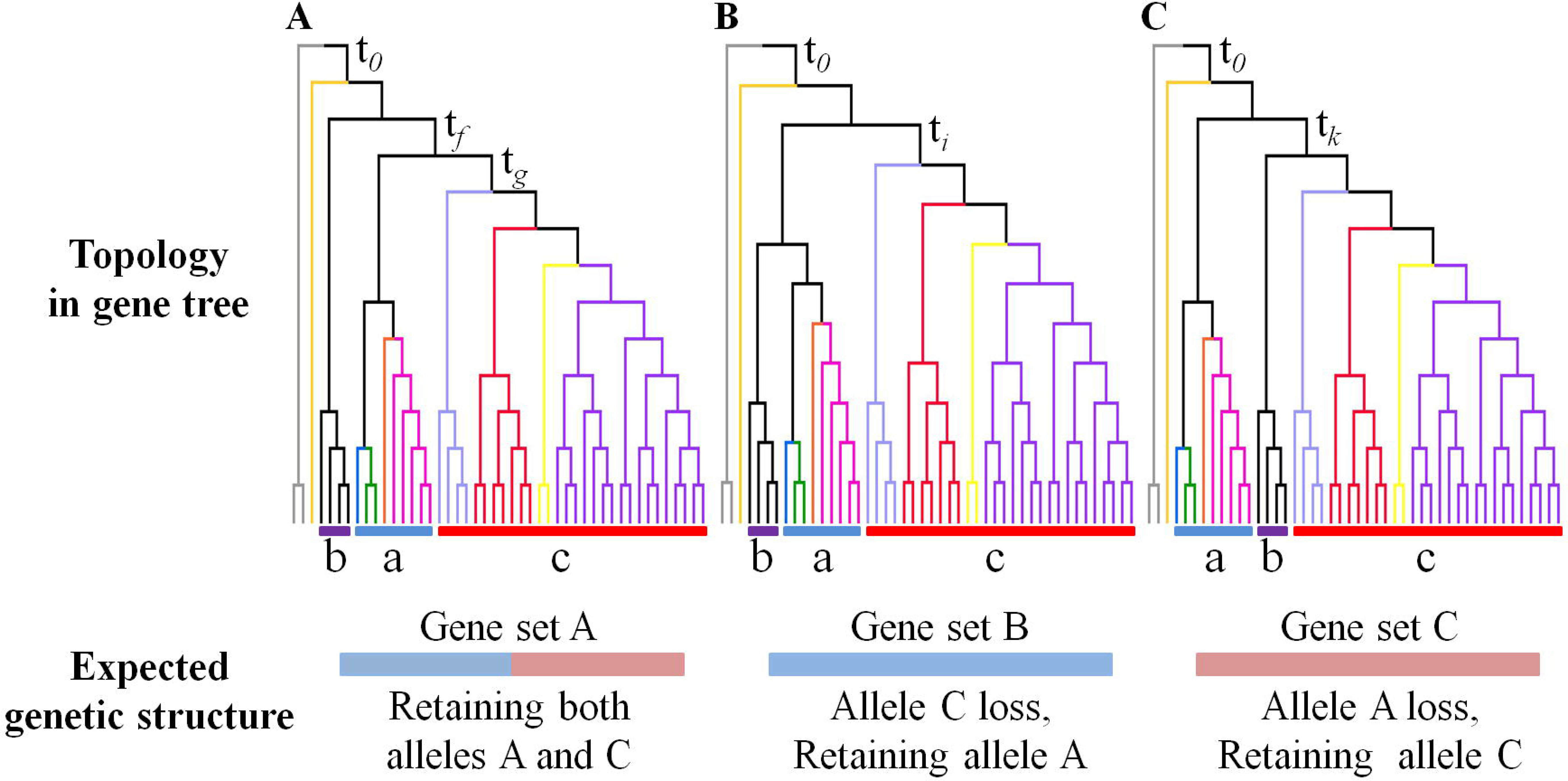
Scheme of the expected selective pressure corresponding to the three position of the *sanguinolenta* group. a, superclade A including four subgenera, b, the *sanguinolenta* group, c, superclade C including subgenus *Stachygynandrum sensu* Weststrand and Korall (2016a). t*_0_*, t*_f_*, t*_g_*, t*_i_*, t*_j_*, and t*_k_*, indicate divergence times.

The divergence times estimated using the two gene sets, in which the *sanguinolenta* group was sister to each superclade, indicated a remarkably consistent time for splits between the *sanguinolenta* group and the two superclades A and C, respectively (Figure S4). This evidence also implies and supports the hybrid origin of the *sanguinolenta* group between the two superclades. Therefore, our molecular dating indicated that the hybridization that created the *sanguinolenta* group occurred during the Early Triassic after the Permian–Triassic extinction (Figure 5A), and the ancient hybridization was supported by the detection of ancient gene flow in the HyDe test (Figure 3), and the inference of hybridization in the PhyloNet result (Figure 4A). Moreover, plastome-based phylogenies showed that the *sanguinolenta* group was sister to the superclade C (Zhang et al., 2020; Chen et al., 2022; Zhang and Zhang, 2022), implying that the common ancestor of the superclade C is probably involved in the hybridization that created the *sanguinolenta* group.

The hybrid origin of the *sanguinolenta* group was also supported by morphological and chromosomal evidence. The leaf morphology was similar to that of the superclade C (subgenus *Stachygynandrum*), while the rhizophores resembled those of the superclade A (four subgenera) (Figure S1; Table S1). Their spores exhibited an intermediate form between the two superclades (Figure S2), and the chromosome number indicated triploidy (Figure S3). The intermediated spore sculptures and triploid chromosomes of the *sanguinolenta* group have also been observed in previous studies (Takamiya, 1993; Zhou et al., 2015b).Consequently, we hypothesized that the enigmatic *sanguinolenta* group was originated by an ancient hybridization between the common ancestors of the two superclades during the Early Triassic after the Permian–Triassic extinction (Figure 5A). Based on the current distribution and habitats of section *Ruperstrae* (superclade A), the *sinensis* group (superclade C), and the *sanguinolenta* group, we speculated that the ancestral distribution of the three groups might overlap in the Induan stage (current northern part of China and the Siberian region) (Figure 5B), where the predicted ancient climate is similar to the current climate with moderately low precipitation (Parrish et al., 1982; Barron and Fawcett, 1995), providing a chance for the ancient hybridization within this family.

It is necessary to discuss how the *sanguinolenta* group has three positions in phylogenetic reconstruction. It has been reported that the species created by an ancient hybridization have three different phylogenetic positions in walnuts (Zhang et al., 2019), and this pattern was also observed in the *sanguinolenta* group (Figure 1). We speculate that three positions of the *sanguinolenta* group may have been induced by selective pressure (Figure 6). For example, the gene showing the topology that formed a sister group with the superclade A retained the allele from the common ancestor of the superclade A, and lost the allele from the other parent (case B in Figure 6). However, the gene showing the topology that formed a sister group to both superclades retained both alleles from the common ancestors of the two superclades, like a random selection over a long time (case A in Figure 6). Thus, the third position, which is not relevant to any parent, might be a signature of the species created by ancient hybridization.

In conclusion, our findings provide evidence that conflicting positions of the *sanguinolenta* group are likely to be caused by an ancient hybridization, and those of the superclade A may be caused by the ILS. We believe that our study not only improves understand the evolutionary history of the Selaginellaceae, and advances our current understanding of the ancient hybridization. Further studies are necessary to reveal the causes of phylogenetic incongruence within the superclade C (subg. *Stachygynandrum*) and demonstrate how the *sanguinolenta* group succeeded in the diploidization to produce spores despite its triploid origin, unlike other triploid bananas and watermelons.

## Materials and Methods

### Taxon sampling and data retrieval

A total of 46 accessions representing 40 species (two *Isoetes* as outgroups) were used for this study (Table S2). Thirty-four accessions of 27 *Selaginella* species were newly sampled and sequenced in this study (Table S2). Among the remaining species, RNA-seq data of seven species were downloaded from 1KP, and those of two species were downloaded from the Sequence Read Archive (SRA) database in NCBI GenBank (Table S2). In addition, whole-genome data of *S. moellendorffii* was used (Banks et al., 2011), and RNA-seq data of two *Isoetes* species from 1KP and NCBI GenBank were downloaded for the outgroup (Table S2). The species sampled in this study covered all six subgenera in the classification of Zhou and Zhang (2015) and six of seven subgenera in that of Weststrand and Korall (2016a) (Table S3).

### RNA sequencing and assembly

For newly sequenced species, total RNA was extracted from mixed tissue samples of stems, leaves, and s porophylls of plants growing in the greenhouse of the Institute of Botany, Chinese Academy of Science s. Paired-end reads of 2×150 were generated by the Illumina HiSeq4000 platform at Novogene Co. Ltd. (Beijing, China). After filtering out reads, clean reads were used for *de novo* assembly using Trinity v. 2. 0.6 with default settings (Grabherr et al., 2011). Redundant nucleotide sequence removal was performed using CD-HIT v4.6 with a threshold of 0.98 (Fu et al., 2012). The remaining transcripts were converted into amino acid sequences using TransDecoder, and the longest peptide for each gene was selected as the potential unigenes.

### Orthologous Genes Selection

To select the orthologous genes, OrthoFinder v.1.1.8 was used for clustering the homologous gene groups (Emms and Kelly, 2015). Among the clustered gene groups, paralogous gene removal was carefully performed in two independent steps. Redundant copies and putative paralogous genes were removed as described previously (Ran et al., 2018). In the first step, using 13 accessions representating 13 different lineages in Selaginellaceae, if each species has at least one transcript, and each subgenus shows monophyly, the homologous gene group was selected for the second step. All data from 46 accessions were used for the second paralogs removal. The two criteria in the first step were also applied in the second step, and the aligned length was considered. When the aligned length of the gene was shorter than 1,000 bp, the gene was excluded for further analyses. Although the species has only one copy of the gene, the possibility of a paralog arising from duplication and loss of the ortholog could not be completely ruled out (see Figure S5 for examples of selected and excluded genes). Therefore, we carefully performed the phylogenetic reconstructions and confirmation of monophyly for all subgenera. For phylogenetic reconstruction of each gene group, sequence alignments were performed using MAFFT (Katoh et al., 2002), and single gene trees were constructed using RAxML v. 8 (Stamatakis, 2006) with 1000 bootstrap replicates and the PROTGAMMAAUTO model. Detailed information on the 347 selected orthologous genes in this study is provided in Table S4.

### Phylogenetic analyses

While selecting orthologous genes with phylogenetic reconstructions, three different positions of the *S. sanguinolenta* group were discovered (A–C in Figure S6). Two of the three positions were previously reported (Figure S6A, C), and one position was newly found in this study (Figure S6B). For further analyses, the selected orthologous genes were divided into three gene sets according to the position of the *S. sanguinolenta* group. Amino acid sequences of each selected orthologous gene were aligned using MAFFT, and the alignments were manually inspected to exclude the sequences with low quality. Phylogenetic analyses were performed using two approaches. For the supermatrix approach, in which all the aligned sequences of orthologous genes concatenate into one matrix, a perl script, catfasta2phyml.pl (https://github.com/nylander/catfasta2phyml), was used to concatenate the aligned orthologous gene sequences. A total of four concatenated supermatrices were prepared, three of which were concatenated sequences of each divided gene set, and one of which was concatenated sequences of all three gene sets. Phylogenetic reconstructions of the four supermatrices were conducted by the ML method using RAxML with 1000 bootstrap replicates and the PROTGAMMAAUTO model. For the supertree approach, single gene trees were constructed using RAxML independently with 1000 bootstrap replicates and the PROTGAMMAAUTO model, and then coalescent-based analyses were performed using ASTRAL (Mirarab et al., 2014). Single gene trees of the three divided gene sets were rooted using Newick Utilities v. 1.7.0 (Junier and Zdobnov, 2010), and coalescent-based analyses were performed for each divided gene set and for all three gene sets together as well. The local posterior probabilities of the species tree were estimated using ASTRAL as node support. We used DensiTree (Bouckaert, 2010) and Dendroscope v. 3.6.3 (Huson and Scornavacca, 2012) to visualize and manipulate the results of phylogenetic analyses.

### Intergroup gene flow and species network analyses

To test intergroup gene flow between clades and to determine the hybrid origin of the *sanguinolenta* group, HyDe was used (Blischak et al., 2018). This tool is similar to the ABBA-BABA test (also known as the *D-* statistic) and has been shown to provide helpful information for detecting hybridization based on phylogenetic invariants (Blischak et al., 2018). To perform HyDe, the concatenated and aligned nucleotides of 444,491 bp in length were used, and all 46 accessions in this study were divided into an outgroup and a triplet of ingroups: two *Isoetes* and *S. selaginoides* as outgroups, species in the superclade composed of the four subgenera as the first ingroup, species in subg. *Stachygynandrum* including the *sinensis* group as the second ingroup, and species in the *sanguinolenta* clade as the third ingroup. If there is no hybridized group, the γ value is 0. A violin plot of 1000 bootstrap replicates was drawn using the violin package in R v.3.5.1 (R Development Core Team, 2015). The species network was inferred using the pseudo-maximum likelihood method (InferNetwork_MPL) implemented in PhyloNet v.3.8.0 (Than et al., 2008). Single gene trees of three different gene sets constructed using RAxML were evenly used for this analysis (A–C in Figure S6). The single-gene trees were rooted using Newick Utilities. Network inference was conducted with settings of 0 to 3 reticulations, specifying the *sanguinolenta* group as hybrid species, and optimizing branch lengths and inheritance probabilities (γ) under the pseudo-likelihood. The inferred species network was displayed by Dendroscope v. 3.6.3.

### Spore and chromosome observations

We examined the surface sculpture of spores using scanning electron microscopy (SEM). Spores were collected from mature strobili of voucher specimens deposited in PE or fresh individuals cultivated in the green house of the Institute of Botany, Chinese Academy of Sciences (IBCAS). Plant materials of *S. vardei* and *S. sanguinolenta* collected from Sichuan, Kangding, and *S. sinensis* collected from Beijing were selected as the representatives of three phylogenetic clades. Megaspores were affixed on a sample stub with doubleLsided adhesive tape directly, while unopened sporangia containing microspores were attached to the stub and smashed with dissecting needle to release microspores. The sample stub was then sputterLcoated with platinum. Observation and photography of spores were conducted with a Hitachi SL4800 (Tokyo, Japan) field emission SEM. For chromosome observation, fresh individuals of *S. sanguinolenta* were used, which were grown in the green house of IBCAS. Vigorous root tips were fixed in Carnoy I solution (3:1 ethanol to glacial acetic acid) at 4 °C for at least three hours after being pre-treated with an ice-water mixture in a dark room for 24 hours. The fixed tips were digested at 37 °C in a combination (1:1) of 2 % cellulase and 2 % pectinase for 60 minutes before staining with an improved carbol-fuchsin solution and being squashed for the observation. The photographs were taken using a Zeiss Axio Imager A1 camera.

### Molecular dating for the ancient hybridization

Currently available methods for molecular dating can only be applied to bifurcating species, but hybridization is not bifurcating. Because our data suggested that the *sanguinolenta* group was originated from hybridization between the common ancestors of each superclade (A and C), we calculated divergence time using gene sets B and C: the *sanguinolenta* group being sister to superclade A in gene set B and sister to superclade C in gene set C, respectively. The divergence time estimation for *Selaginella* species was performed using penalized likelihood (PL) implemented in r8s v.1.81 (Sanderson, 2003) and MCMCtree implemented in PAML v.4.9 (Yang, 2007). In both methods, the same two fossils were used as calibration points. The split between Isoetaceae and Selaginellaceae was calibrated at 370 Ma based on the *Lepidosigillaria* reported in the Late Devonian (Kenrick and Crane, 1997). Klaus et al. (2017) used isophyllous *Selaginella resimus* to fix the split between the *sanguinolenta* group and all other *Selaginella* species at 330–350 Ma based on the topology A in Figure S5 as the backbone, but the *sanguinolenta* group is anisophyllous. Moreover, the conflicting positions of the *sanguinolenta* group revealed here indicate the inappropriation putting the fossil of *S. resimus* as calibration point. The fossil of *S. suissei* found in the Late Carboniferous (ca. 304 Ma) (Zeiller, 1906) was used to calibrate the split between subg. *Selaginella* (*S. selaginoides*) and the remaining *Selaginella* species. By the PL method, the ML trees based on the four concatenated matrices resulted from RAxML were used as the input. Prior to estimating the divergence time, a cross-validation test was performed to determine the best smoothing value for each tree. With the best smoothing values, the divergence times were estimated with the TN and default settings of the other parameters. For the MCMCtree method, the rough substitution rates were estimated using the CODEML in PAML, and then the gradient (*g*) and Hessian (*H*) were estimated using the MCMCtree in PAML for each dataset. Finally, the divergence times were estimated using the calculated rgene_gamma and sigma2_gamma. The MCMC chain was analyzed for 10,000,000 generations with a sample frequency of 50 and a burn-in phase of 1,000,000 generations. We used Tracer v.1.7 (Rambaut et al., 2018) to check that the effective sample size was greater than 200. The estimated divergence time trees were displayed by FigTree v.1.4.3. We compared the estimated divergence times for splits between the *sanguinolenta* group and the two superclades inferred from the two gene sets independently and then visualized the divergence time inferred from gene set C as the backbone of molecular dating for the hybridization occurrence.

## Data availability

All relevant data can be found within the manuscript and its supporting information. Raw RNA sequencing data generated in this study are available at the Sequence Read Archive of NCBI GenBank, under BioProject PRJNA945812. The voucher specimens were deposited in the Chinese National Herbarium (PE), and other data and material provided in this manuscript are available from the corresponding author upon reasonable request.

## Acknowledgments

This study was supported by grants from the National Natural Science Foundation of China (Grant Numbers 32170233 and 32270248), and a grant from the Youth Innovation Promotion Association CAS (Grant Number 2021075).

## Authors’ contributions

QPX and XCZ contributed to the study conception and design. Material preparation was performed by XCZ. Experiments and data analyses were performed by JSK. Data collection and interpretation were performed by JSK and JGY. The first draft of the manuscript was written by JSK, QPX, and XCZ.

## Supporting Information

**Text S1. Previously reported phylogenetic incongruences of two enigmatic groups in Selaginellaceae.**

**Text S2. Orthologous gene selection and paralogous gene removal.**

**Figure S1. Rhizophores on dorsal side in the Selaginella *sanguinolenta* group.** (A and B) Rhizophores of *Selaginella rossii*. (C) Rhizophore of *Selaginella sanguinolenta*. The white dashed circle presents the rhizophore on the dorsal side.

**Figure S2. Megaspores and microspores of three representative *Selaginella* species.** *S. vardei* belongs to subgenus *Rupestrae* (superclade A), *S. sanguinolenta* represents the *sanguinolenta* group, and *S. sinensis* belongs to subgenus *Stachygynandrum* (superclade C).

**Figure S3. Thirty chromosomes** of *Selaginella sanguinolenta*. Somatic chromosomes of metaphase in *S. sanguinolenta*.

**Figure S4. Divergence time estimations for the Selaginellaceae using two different methods.** The red stars indicate fossil calibration points. Divergence times were estimated using gene set B (left) and gene set C (right). (A) Divergence time estimated by Penalized likelihood (PL) method. (B) Divergence time estimated by MCMCtree method.

**Figure S5. Conceptual differences between hybridization and incomplete lineage sorting.** Both concepts of hybridization and incomplete lineage sorting were adopted from Sang and Zhong (2000) and Pamilo and Nei (1988). Representative species trees, where “a”, “b”, and “c” are ingroup species and o is an outgroup species. t*_0_*indicate time of speciation between ingroup and outgroup. t*_f_*, t*_g_*, t*_i_*, t*_j_*, t*_k_*, and t*_m_* indicate divergence times, and t*_h1_* and t*_h2_* indicate time when the lineages that hybridized to give rise to “b” diverged from “a” and “c”. Representative species trees are given in (a, b). The two different gene trees are given in (c, d, e, and f). Expected gene-wise frequencies in the genome shown as near trees (b–f).

**Figure S6. Examples of selected and excluded single gene trees in this study.** *Out*: outgroup, *Se*: subg. *Selaginella*, *Er*: subg. *Ericetorum*, *Gy*: subg. *Gymnogynum*, *Le*: subg. *Lepidophyllae*, *Ru*: subg. *Rupestrae*, *St*: *Stachygynandrum* group, *He*: *heterostachys* group, *Pu*: the *pulvinata* group, *Sa*: the *sanguinolenta* group, *Si*: the *sinensis* group. He, Pu, Sa, and Si belong to subgenus *Stachygynandrum* in the latest classification (Weststrand and Korall, 2016a). Single gene tree showing monophyletic for each lineage was selected for further analyses (A-C). Single gene tree showing nonmonophyletic in some lineage was excluded (D-F).

**Table S1. Comparison of morphological and chromosomal characters from each group within Selaginellaceae.**

**Table S2. List of 46 accessions and their information used in this study.**

**Table S3. Comparison of three representative infrageneric classifications in the genus *Selaginella*.**

**Table S4. Functional information of the nuclear genes used for reconstructing phylogenies in this study.**

