## Supplemental figures 1-6 for "The earliest allopolyploidization in tracheophytes revealed by phylotranscriptomics and morphology of Selaginellaceae"

**Text S1.** **Previously reported phylogenetic incongruences of two enigmatic groups in Selaginellaceae.**

Phylogenetic studies have revealed two enigmatic groups, *Selaginella sanguinolenta* and *S. sinensis* groups in Selaginellaceae. The *S. sanguinolenta* group was sister to all rhizophoric *Selaginella* species, except for subgenus *Selaginella* (*S. selaginoides* and *S. deflexa*) based on the plastid *rbcL* and nuclear ribosomal ITS region (Zhou et al., 2015a). In another study, the sister relationship between the *sanguinolenta* group and other rhizophoric clade was supported by the Bayesian Inference (BI) tree based on the plastid *rbcL* and nuclear *SQD1* genes, while the other position of the *sanguinolenta* group, sister to subgenus *Stachygynandrum* in the BI tree, was discovered based on nuclear *pgiC* gene (Weststrand and Korall, 2016b). Similarly, the *sinensis* group was located in the most basal clade which was sister to all *Selaginella* species, even including subgenus *Selaginella*, in the Maximum Parsimony (MP) tree based on plastid *rbcL* sequences, but it was resolved as sister to subgenus *Stachygynandrum* in the BI tree based on *rbcL* and in both MP and BI trees based on 26S rDNA sequences (Korall and Kenrick, 2004). Another study indicated that the *sinensis* group was sister to the other five subgenera (*Rupestae*, *Lepidophyllae*, *Gymnogynum*, *Exaltatae*, *Ericetorum*) in the BI tree using *rbcL* sequences, but they found an incongruence between two single-copy nuclear gene trees (*pgiC* and *SQD1*), like the incongruence of the *sanguinolenta* group (Weststrand and Korall, 2016b). Phylogenetic positions of these two enigmatic groups were still not resolved even though plastid genome (plastome) and mitochondrial genome (mitogenome) data were used for the phylogenetic reconstruction. Plastome-based phylogenies from indicated that the *sanguinolenta* group was located within the subgenus *Stachygynandrum*, but their positions differed in the two plastome-based phylogenies (Zhang et al., 2020; Zhou et al., 2022). However, the mitogenome-based phylogeny indicated a sister relationship between the *sanguinolenta* group and other *Selaginella* species (including superclade A and C in this study), except for subgenus *Selaginella* (Tang et al., 2023). Like the *sanguinolenta* group, the *sinensis* group was located in the most basal clade, and sister to all other *Selaginella* species in the plastome-based phylogeny (Zhou et al., 2022), while the group was sister to the clade of the *Heterostachys* and *pulvinata* groups in the mitogenome-based phylogeny (Tang et al., 2023).

**Text S2. Orthologous gene selection and paralogous gene removal.**

More than 3,000 single/low copy nuclear gene groups were used at the beginning of orthologous gene selection, and 347 nuclear genes were eventually selected for the phylogenetic reconstruction. Using 13 representative species, the first removal was performed with two criteria; all 13 species must have at least one transcript, and each group should be monophyletic in each tree constructed by ML method. After the first removal, 1025 putative orthologous genes remained. For the second round, all 46 accessions were included (Table S2), and three criteria were applied that all species have at least one transcript, showing monophyletic for each group in the gene tree, and the aligned length of the gene should be over 1,000 bp. The monophyly of each group was confirmed manually. After all, a total of 347 orthologous genes remained, and were used for further analyses. However, while confirming the monophyly for each group, we found that the *sanguinolenta* group was located at three different positions, with two positions reported previously, and one newly found in this study (Figure 1). According to the positions of the *sanguinolenta* group, the three different gene groups are referred to as gene sets A, B, and C, respectively. Gene set A consists of 130 orthologous genes and the *sanguinolenta* group is sister to all rhizophoric *Selaginella* species, including two superclades (Figure 1A), gene set B consists of 81 orthologous genes and the *sanguinolenta* group is sister to the clade composed of the other four subgenera (Figure 1B), and gene set C consists of 136 orthologous genes and the *sanguinolenta* group is sister to subgenus *Stachygynandrum* (Figure 1C). Within each gene set, orthologous genes were not related to the functional roles (Table S4), indicating that the constructing of different topologies in each gene set is not relevant to gene functions. The information on 347 selected orthologous genes in this study was described in Table S4.

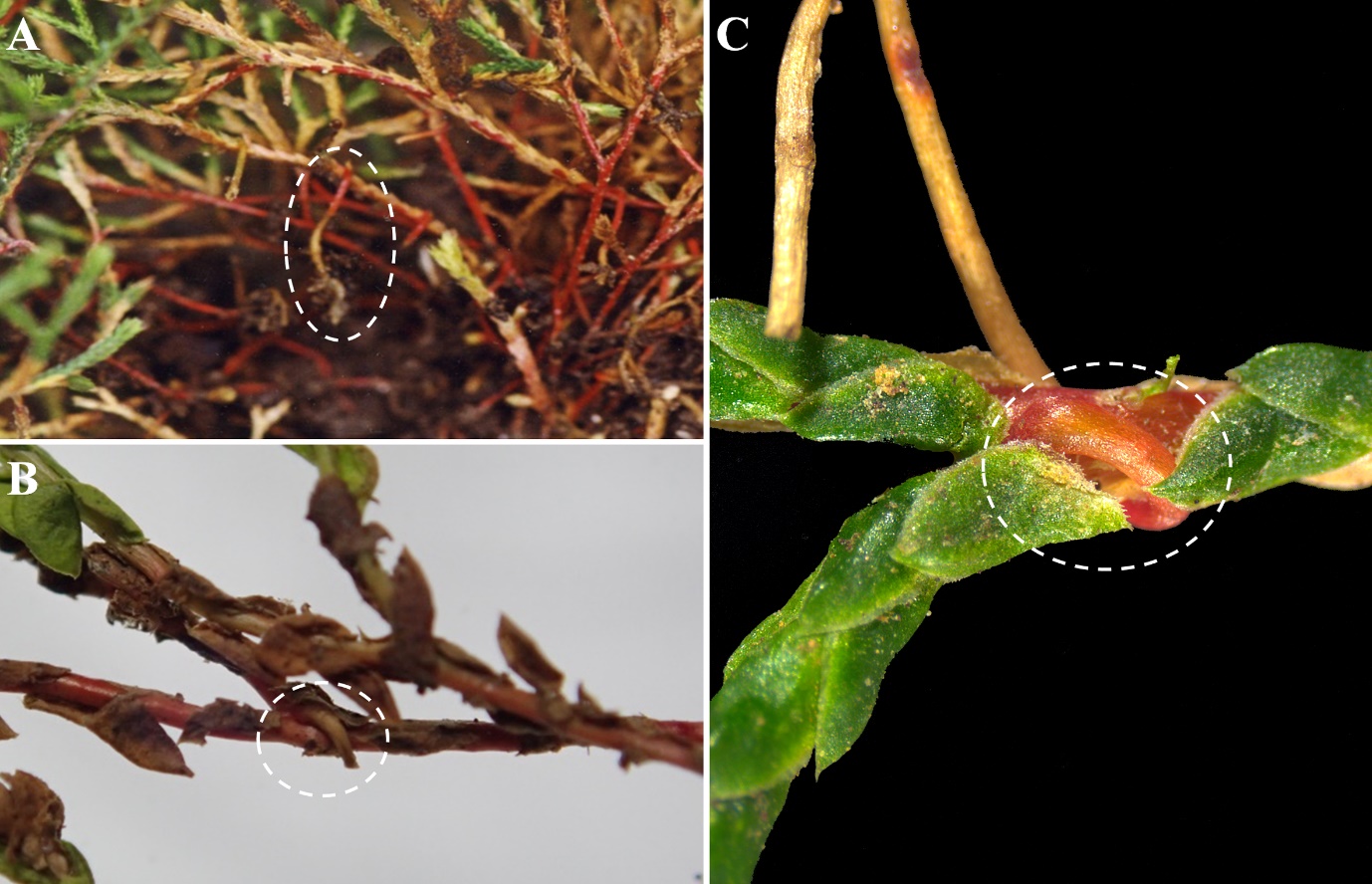

**Figure S1. Rhizophores on dorsal side in the *Selaginella sanguinolenta* group.** (A and B) Rhizophores of *Selaginella rossii*. (C) Rhizophore of *Selaginella sanguinolenta*. The white dashed circle presents the rhizophore on the dorsal side.

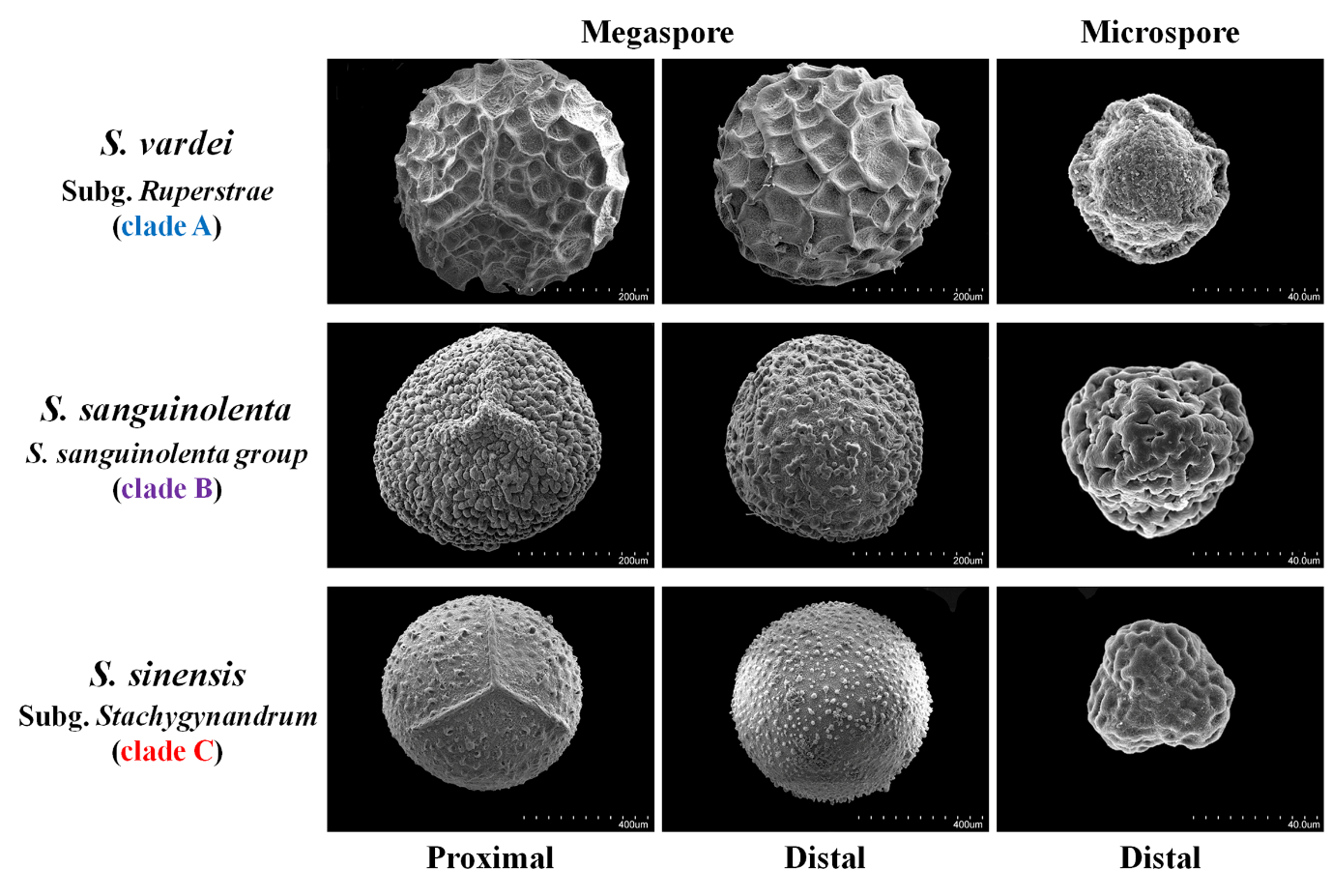

**Figure S1. Megaspores and microspores of three representative *Selaginella* species.** *S. vardei* belongs to subgenus *Rupestrae* (superclade A), *S. sanguinolenta* represents the *sanguinolenta* group, and *S. sinensis* belongs to subgenus *Stachygynandrum* (superclade C).

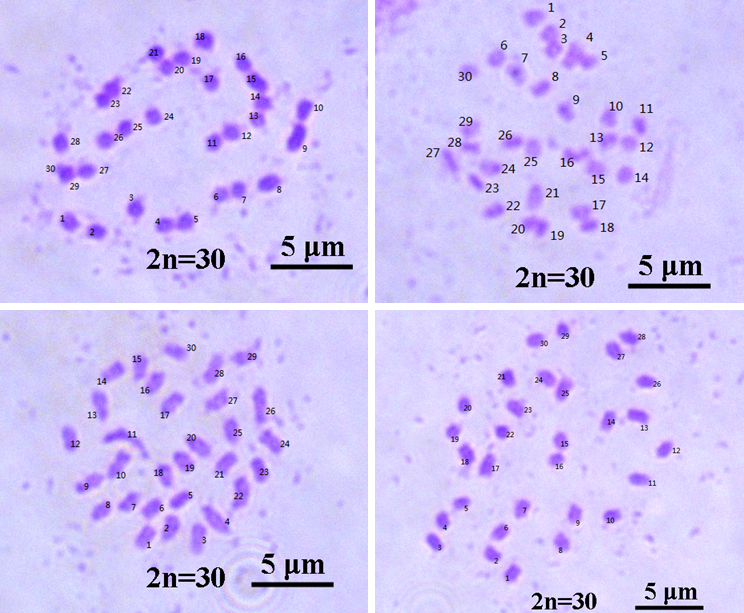

**Figure S3. Thirty chromosomes of *Selaginella sanguinolenta*.** Somatic chromosomes of metaphase in *S. sanguinolenta*.

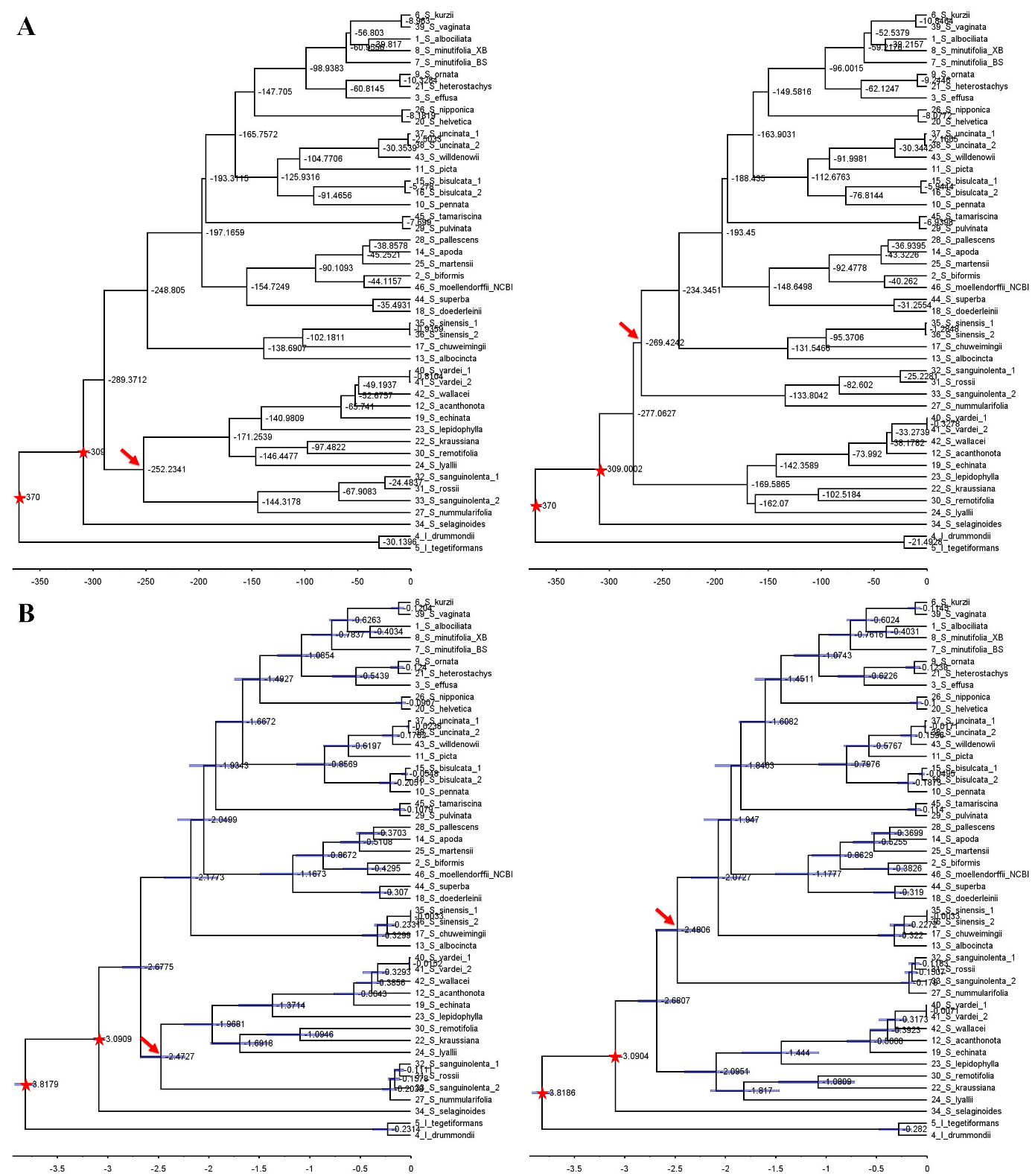

**Figure S4. Divergence time estimations for the Selaginellaceae using two different methods.** The red stars indicate fossil calibration points. Divergence times were estimated using gene set B (left) and gene set C (right). (A) Divergence time estimated by Penalized likelihood (PL) method. (B) Divergence time estimated by MCMCtree method.

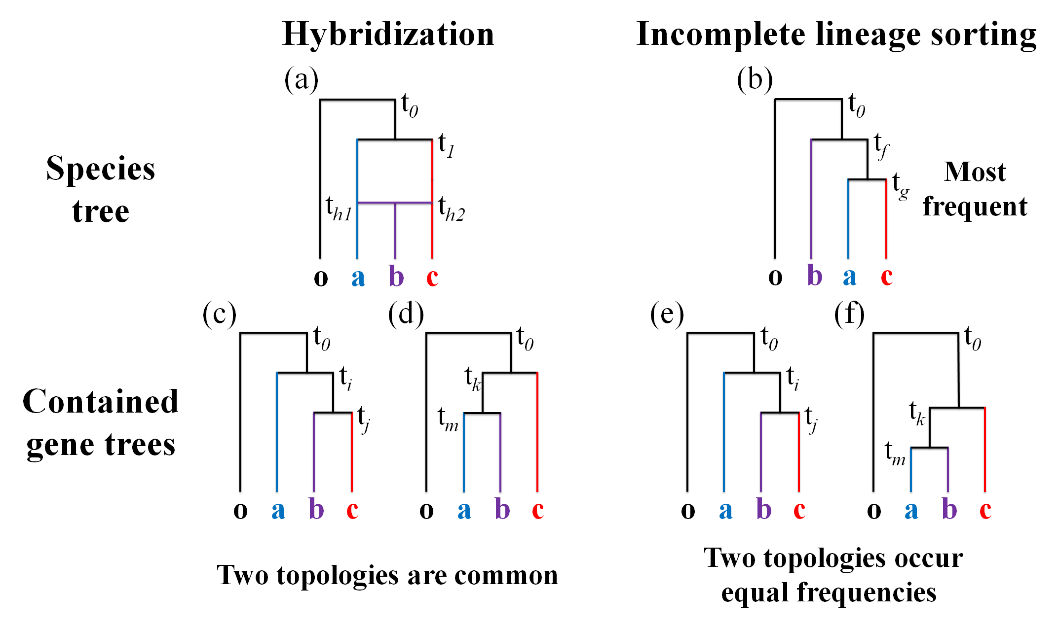

**Figure S5. Conceptual differences between hybridization and incomplete lineage sorting.** Both concepts of hybridization and incomplete lineage sorting were adopted from Sang and Zhong (2000) and Pamilo and Nei (1988). Representative species trees, where “a”, “b”, and “c” are ingroup species and “o” is an outgroup species. t*_0_* indicates the speciation time between the ingroup and outgroup. t*_f_*, t*_g_*, t*_i_*, t*_j_*, t*_k_*, and t*_m_* indicate divergence times, and t*_h1_* and t*_h2_* indicate the hybridization time giving rise to “b” diverged from “a” and “c”. Representative species trees are given in (a, b). The two different gene trees are given in (c, d, e, and f). Expected gene-wise frequencies in the genome are shown near trees (b–f).

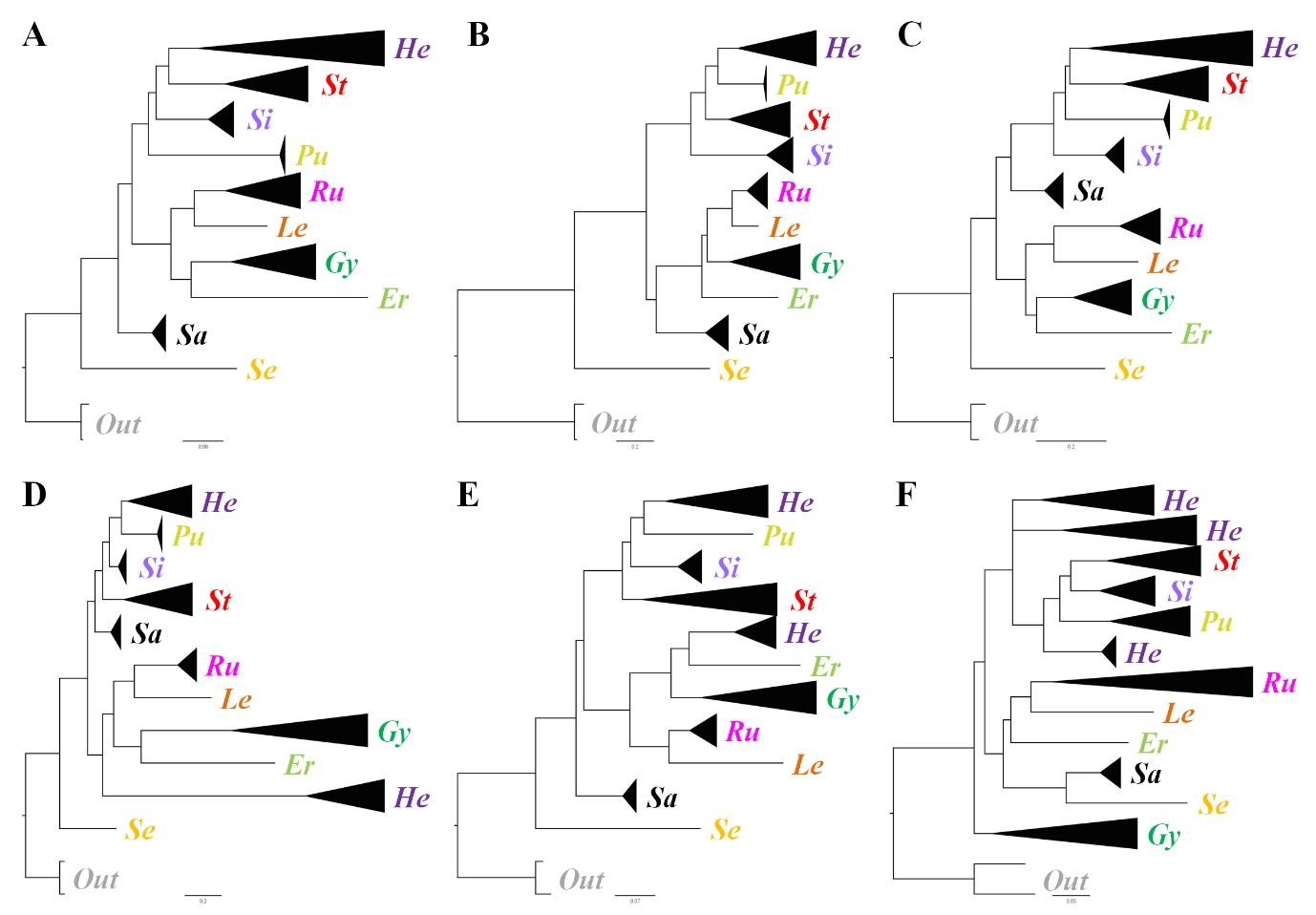

**Figure S6. Examples of selected and excluded single gene trees in this study.** *Out*: outgroup, *Se*: subg. *Selaginella*, *Er*: subg. *Ericetorum*, *Gy*: subg. *Gymnogynum*, *Le*: subg. *Lepidophyllae*, *Ru*: subg. *Rupestrae*, *St*: the *Stachygynandrum* group, *He*: the *Heterostachys* group, *Pu*: the *pulvinata* group, *Sa*: the *sanguinolenta* group, *Si*: the *sinensis* group. He, Pu, Sa, and Si belong to subg. *Stachygynandrum* *sensu* Weststrand and Korall (2016a). The single gene trees showing monophyletic for each lineage were selected for further analyses (A-C). The single gene trees showing nonmonophyletic in some lineages were excluded (D-F).
